# Unsupervised encoding selection through ensemble pruning for biomedical classification

**DOI:** 10.1101/2022.02.06.479282

**Authors:** Sebastian Spänig, Alexander Michel, Dominik Heider

## Abstract

**Background:** Owing to the rising levels of multi-resistant pathogens, antimicrobial peptides, an alternative strategy to classic antibiotics, got more attention. A crucial part is thereby the costly identification and validation. With the ever-growing amount of annotated peptides, researchers leverage artificial intelligence to circumvent the cumbersome, wet-lab-based identification and automate the detection of promising candidates. However, the prediction of a peptide’s function is not limited to antimicrobial efficiency. To date, multiple studies successfully classified additional properties, e.g., antiviral or cell-penetrating effects. In this light, ensemble classifiers are employed aiming to further improve the prediction. Although we recently presented a workflow to significantly diminish the initial encoding choice, an entire unsupervised encoding selection, considering various machine learning models, is still lacking.

**Results:** We developed a workflow, automatically selecting encodings and generating classifier ensembles by employing sophisticated pruning methods. We observed that the Pareto frontier pruning is a good method to create encoding ensembles for the datasets at hand. In addition, encodings combined with the Decision Tree classifier as the base model are often superior. However, our results also demonstrate that none of the ensemble building techniques is outstanding for all datasets.

**Conclusion:** The workflow conducts multiple pruning methods to evaluate ensemble classifiers composed from a wide range of peptide encodings and base models. Consequently, researchers can use the workflow for unsupervised encoding selection and ensemble creation. Ultimately, the extensible workflow can be used as a plugin for the PEPTIDE REACToR, further establishing it as a versatile tool in the domain.

## Background

Multi-resistant pathogens are a major threat for modern society [1]. In the last decades, a rising number of bacterial species developed mechanisms to elude efficiency to widely used antibiotics [1]. The importance of developing and implementing alternative strategies is further underpinned by a recent study, which detected a certain baseline resistance in European freshwater lakes [2]. The study confirmed resistance specifically against four critical drug classes in human and veterinary health in freshwater, which is typically considered as a pathogen-free environment [2]. Moreover, already concerning levels of antibiotic resistance in Indian and Chinese lakes emphasize the requirement of alternative biocides [3, 4]. One promising approach to replace or even support common antibiotics refers to the deployment of peptides with antimicrobial efficiency [5]. However, identifying and validating active peptides requires intensive, hence, costly and time-consuming wet-lab work. Thus, in the pre-artificial intelligence (AI) era, the manual classification and verification of antimicrobial peptides (AMPs) engaged researchers. Although the *in vitro* confirmation of activity is still necessary, the application of AI, i.e., in particular machine learning (ML) algorithms, simplifies the identification process drastically and pushed several AMPs to the second or third phase of clinical trials [6]. In addition, online databases provide access to thousands of annotated sequences and pave the way automatic peptide design and classification [7]. For instance, Chung *et al*. (2019) developed a method, which demonstrated good performance on classifying AMPs using a two-step approach, which first predicts efficiency, and afterward the precise target activity [8]. Another study employed a variational autoencoder to encode AMPs, mapped the probability of being active to a latent space, and predicted novel AMPs [9]. Fingerhut *et al*. (2020) introduced an algorithm to detect AMPs from genomic data [10]. For more information on computational approaches for AMP classification, we refer to the recent review of Aronica *et al*. (2021) [11].

However, the prediction of amino acid sequence features is not limited to AMPs. In the literature, one can find various applications, e.g., in oncology for predicting anticancer peptides [12], in pharmacology for the discovery and application of cell-penetrating peptides as transporters for molecules [13], or in immunotherapy, for classifying of pro- or antiinflammatory peptides [14, 15]. Other applications include antiviral peptides [16], or peptides with hemolytic [17] or neuro transmitting activity [18].

Unequivocally, the success of ML methods for the prediction of AMPs was enabled by the development and advances of peptide encodings. Encodings are algorithms mapping the amino acid sequences of different lengths to numerical vectors of an equal length, hence, fulfilling the requirement of many ML algorithms [19]. Moreover, peptides or proteins can be described by their primary structure, i.e., the amino acid sequence, and the aggregation in higher dimensions, denoted as the secondary or tertiary structure. Encodings derived from the primary structure are known as sequence-, and encodings describing a higher-order folding are structure-based encodings. To date, a large number of sequence- and structure-based encodings have been introduced and employed in various studies [19]. A significant amount of encodings has been recently acknowledged by another study, specifically benchmarking these by considering multiple biomedical applications [20]. It turned out that most encodings show acceptable performance, partly also beyond single biomedical domains [20]. In addition, Spänig *et al*. (2021) developed a workflow, which can dramatically reduce the number of initial encodings [20]. However, encoding selection is still challenging, and user-friendly approaches are required.

Furthermore, hyperparameter optimization is additionally aggravated by the model choice. Albeit Support Vector Machines (SVM) and Random Forests (RF) are widely employed in peptide classification [11], the variety of models used in a broad range of studies is large. For instance, Khatun *et al*. (2020) utilized several ML algorithms, including Näive Bayes, AdaBoost, and a fusion-based ensemble for the prediction of proinflammatory peptides [21]. The fusion-based model outperformed the other ML models significantly for this task [21]. Plisson *et al*. (2020) employed Decision Trees (DT) and Gradient Boosting (GB), among others, to classify non-hemolytic peptides and demonstrated that the GB ensemble has superior performance [22]. In contrast, Timmons *et al*. (2020) used Artificial Neural Networks to characterize therapeutic peptides with hemolytic activity [23]. Singh *et al*. (2021) compared several base classifiers, e.g., Linear Discriminant Analysis and ensemble methods, e.g., GB and Extra Trees to detect AMPs [24]. They demonstrated that the GB performed best [24]. These studies clearly show that ensemble classifiers typically show superior performance than single classifiers, owing to the fact the fact that they can compensate for weaknesses of single encodings and base classifiers [25].

Recently, Chen *et al*. (2021) introduced a comprehensive tool, which allows less programming experienced researchers to simply select encodings and base or ensemble classifiers through a graphical user interface, allowing easy access to the underlying algorithms [26]. Nevertheless, the approach assumes that the user selects proper settings for the parameterized encodings, which has been previously shown to affect the classification process significantly [20]. Moreover, the encoding selection is independent of the classifier settings, meaning that the tool can set up the classifier automatically; however, the encoding selection is not part of it. Thus, it remains a challenge to pick good encodings and classifiers for a biomedical classification task at hand. To this end, we assessed unsupervised encoding selection and the performance and diversity of multiple ensemble methods. We added different overproduce-and-select techniques for ensemble pruning, facilitating an automatic ensemble generation. In addition, we utilized Decision Trees, Logistic Regression, and Näive Bayes as base classifiers, owing to their prevalence in the field of biomedical classification [11, 19, 27].

Besides demonstrating the benefit of an unsupervised encoding selection, we also examined how the RF performs as a base and ensemble classifier. Specifically, we examined whether the RF, also an ensemble method, is performance-wise already saturated or whether a subsequent fusion can improve the final predictions. Fusion of RFs has been shown in other studies to improve overall performance, e.g., for HIV tropism predictions [28, 29]. All in all, we complement our recent large-scale study on peptide encodings [20] with an automatic encoding selection and a performance analysis of multiple base and ensemble classifiers. Ultimately, the present research bridges the gap between many peptide encodings and available machine learning models.

## Results

We developed an end-to-end workflow, which automatically generates and assesses classifier ensembles using different pruning methods and a variety of encoded datasets from multiple biomedical domains (see Table 5). Researchers can easily extend the workflow with different base and ensemble classifiers, pruning methods, and encoded datasets. The results can be reviewed using the provided visualizations, and the performance is further revised using multiple statistics. We demonstrate that the Pareto frontier pruning is a valuable technique to generate efficient classifier ensembles. However, the utilized base classifiers show comparable performance. We address the results in more detail in the following. We use the example of the avp amppred dataset throughout the manuscript. The results for the remaining datasets can be found in the supplement. Moreover, the code is publicly available at https://github.com/spaenigs/ensemble-performance. The workflow produces interactive versions of all charts (see supplements).

### Pruning methods

All pruning methods generate ensembles, i.e., combined encodings, superior to the single classifiers (see Tables 1 and 2). In the case of the Pareto frontier (pfront) pruning, which is mainly ranked among the best pruning methods, we observe a significant (*p ≤* 0.001) performance improvement compared to the the single classifiers. We also observed that the pfront pruning generates larger ensembles than the convex hull (chull) pruning, which can be visually verified in Fig. 4 (red line). Notably, including the Random Forest (RF) classifier (see Table 1, pfront) does not, or very slightly, affect the ensemble performance without RF (see Table 2), although the the single classifier performance is improved with the RF included (see Table 1). Consequently, the RF increases the overall performance of the ensembles generated by the best encodings pruning.

**Table 1.**
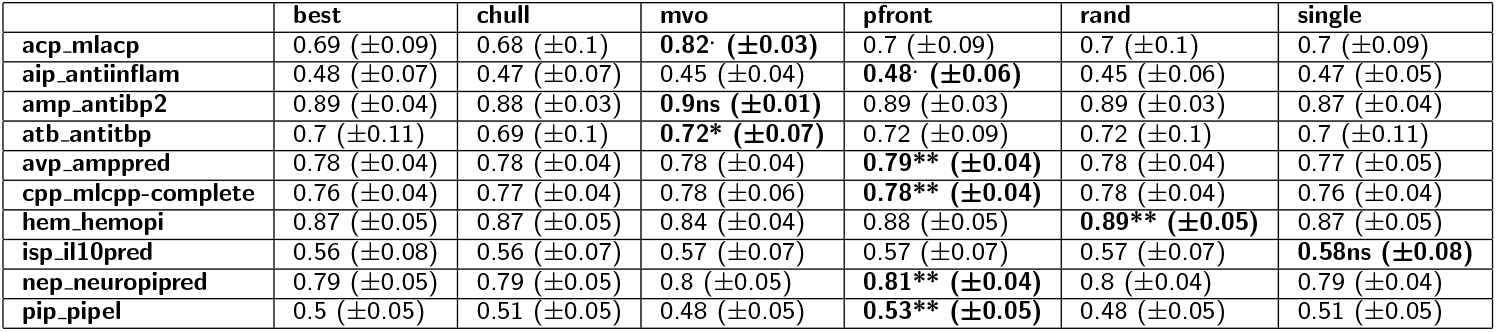
The table shows the performance comparison (including RF) of classifier ensembles derived from different pruning methods, i.e., best, chull, mvo, pfront, rand, and the single best classifier. We used the best ensemble/base model combination, respectively. Numbers refer to the mean performance of a 100-fold Monte Carlo cross-validation. Standard deviation (SD) is added in brackets. Mean and SD are rounded to 2 decimal places. The top base/ensemble classifier combination is always used (see Fig. 2). Classifier ensembles are significantly better than the single best classifiers. In particular, except for one case, the Pareto frontier pruning (pfront) generates the best ensembles. Significance levels are as follows: ** *p ≤* 0.001, * *p ≤* 0.01, and. *p ≤* 0.05. Refer to Table 1 for the reported MCC values from the original studies.

**Table 2.**
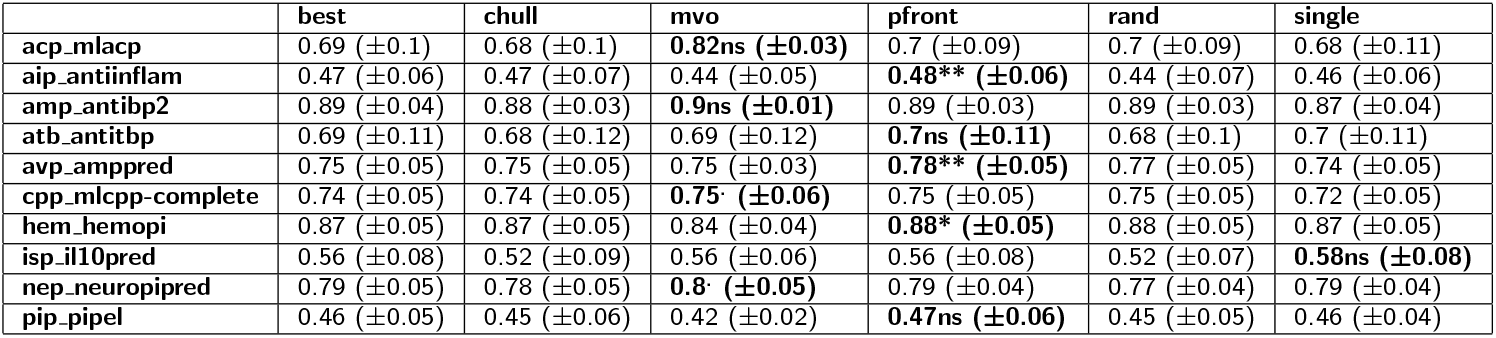
The table shows the performance comparison (excluding RF) of classifier ensembles derived from different pruning methods and the single best classifier. See Table 1 for more details.

**Table 3.** Employed datasets in this study. The function refers to the positive class, i.e., sequences of class + possess the respective function. The stated MCC refers to the performance reported in the original study. See Reference or [20] for more details.

|  | Function | MCC | Size (+,-) | Reference |
| --- | --- | --- | --- | --- |
| acp_mlacp | Anti-cancer | 0.698 | 581 (185,396) | [12] |
| aip_antiinflam | Anti-inflammatory | 0.45 | 2124 (863,1261) | [14] |
| amp_antibp2 | Anti-microbial | 0.84 | 1975 (981,994) | [30] |
| atb_antitbp | Anti-tubercular | 0.52 | 492 (246,246) | [31] |
| avp_amppred | Anti-viral | 0.8 | 1476 (738,738) | [16] |
| cpp_mlcpp | Cell-penetrating | 0.793 | 1901 (737,1164) | [13] |
| hem_hemopi | Hemolytic | 0.52 | 1013 (522,461) | [17] |
| isp_il10pred | Immunosuppressive | 0.59 | 1242 (394,848) | [32] |
| nep_neuropipred | Neuropeptides | 0.67 | 1750 (875,875) | [18] |
| pip_pipel | Pro-inflammatory | 0.454 | 3228 (833,2395) | [15] |

The multi-verse optimization (MVO) suffers from high computational demand, i.e., a long pruning time, however, demonstrates at least comparable performance (see Table 1 and 2). The fitness of the models increases rapidly in the first generations (see Fig. 1). However, towards the last generations the curve flattens. The RF-based models show already higher fitness in the beginning, only increasing slightly in the course of the generations. The robustness of the RF classifier is also highlighed by the low standard deviation across the folds (see Fig. 1).

**Figure 1.**
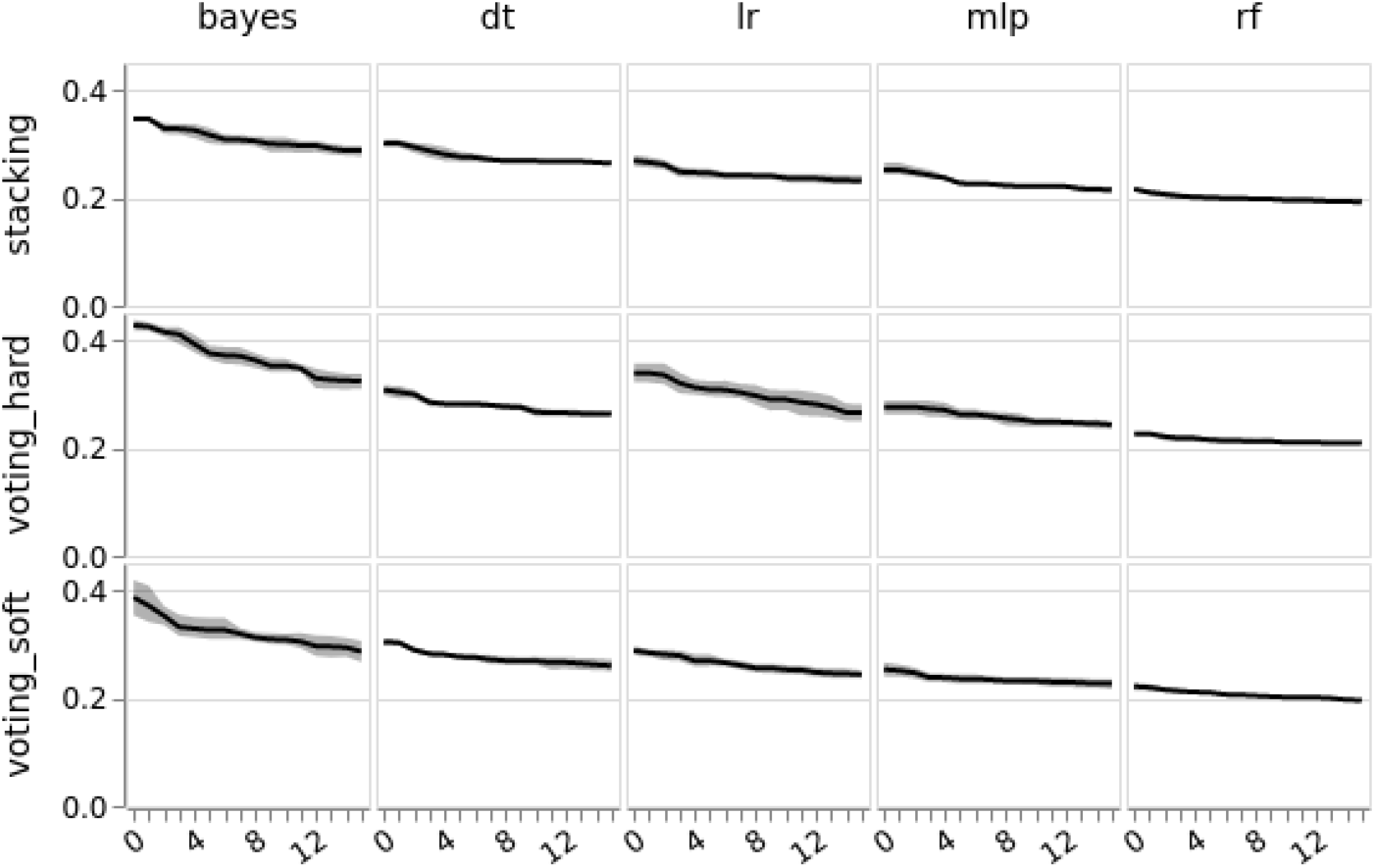
Comparison of the fitness and generations. The x-axis depicts the number of generations and the y-axis the fitness (1 *− MCC*). The respective base classifiers are in the columns and the rows depict the ensemble methods. The light gray area is standard deviation across 5 folds. The plot shows the example of the avp amppred dataset.

### Ensemble classifiers

The ensemble performance mainly depends on the pruning and the choice of the base classifiers; hence, using individual encodings. The performance between the best, random (rand), and chull pruning is insignificant, which stresses the effectiveness of the Pareto frontier (pfront) pruning (see Fig. 2). Furthermore, no significant difference can be observed for ensembles with the same base classifiers, e.g., the RF or Decision Tree (DT). Thus, the fusion method impacts the overall performance only slightly. However, various base classifiers result in significant different ensembles, i.e., employing, for instance, the RF, generates significantly different ensembles compared to the application of other base classifiers (see Fig. 2).

**Figure 2.**
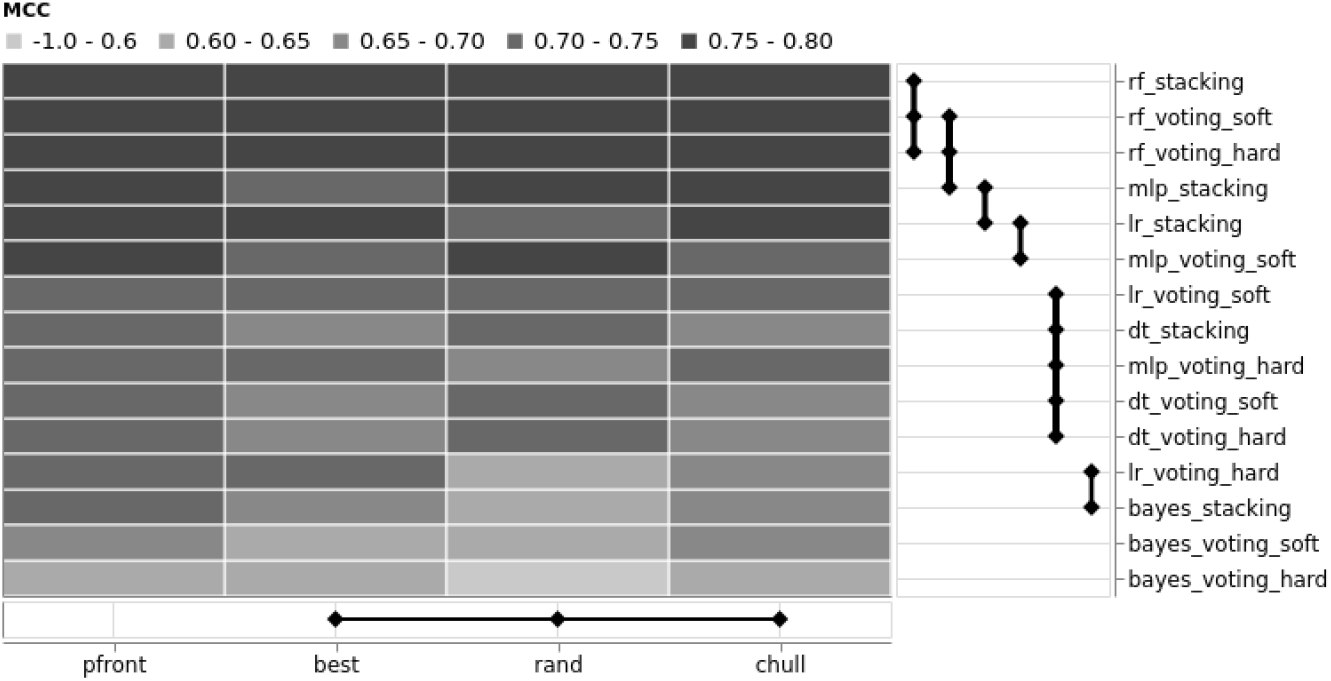
The XCD chart shows the difference between pruning methods (x-axis) and ensembles (y-axis). Entities connected with a bold line are not significant different. The higher the performance, the darker the color. The plot shows the example of the avp amppred dataset.

Moreover, we observed that the Näive Bayes (NB), the Logistic Regression (LR), and the Multi-Layer Perceptron (MLP) classifiers result in ensembles with higher variance (see Fig. 4). In contrast, the area covered by RF and DT models is more compact. Therefore, the variables, i.e., diversity and the pairwise error, are revised by a multivariate analysis of variance (MANOVA), which revealed a significant difference (*p <* 0.001). A separate examination of the variables utilizing variance analysis (ANOVA) followed by a post-hoc analysis using Tukey’s HSD, demonstrates that all variables are significantly different (*p <* 0.001). Finally, we conducted an ANOVA on the particular area values, which disproves the initial observation, i.e., all areas are significantly different (*p <* 0.001). However, considering the average values for all datasets, the DT and RF are commonly ranked as the base classifiers with low variance (see Table 4).

**Table 4.**
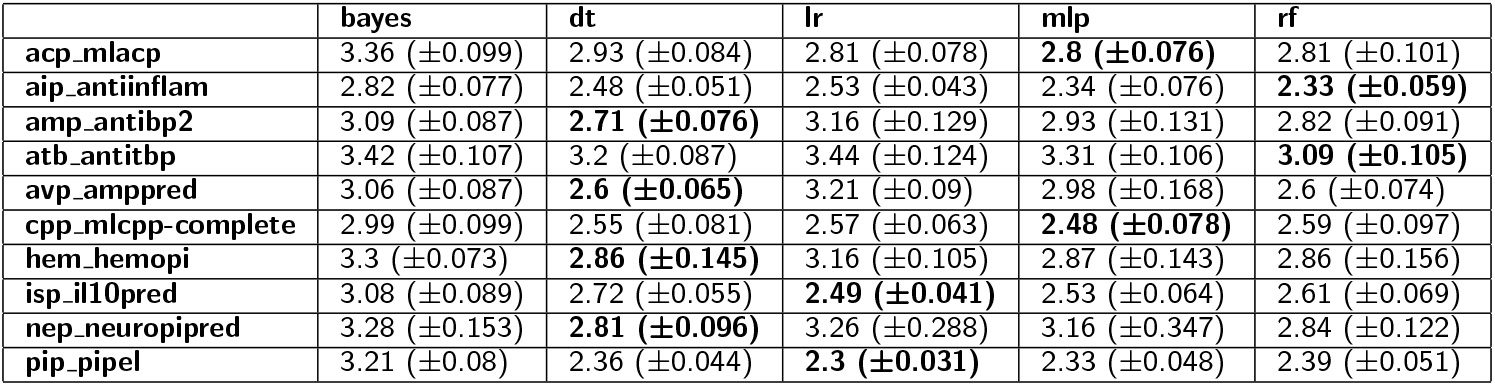
The table lists the average area (*±* SD) covered by the base classifiers across the 100-fold Monte Carlo cross-validation. The lowest area per dataset is highlighted in bold. The DT classifier has the lowest area for most of the datasets, i.e., the predictions are more stable. Refer to Fig. 4 (bottom) for the example showing the avp amppred dataset.

### Single classifiers

In general, the performance of the individual classifiers (single) is lower compared to the classifier ensembles (see Fig. 3). In addition, we noticed that the RF is relatively saturated, i.e., using the RF as a single classifier and as a base model for ensembles does not have a significant effect on performance improvement. The low-performance variance is in line with the observation that weak models benefit most from ensemble learning; however, RFs are ensemble models [33, 34]. In contrast, the performance of other single classifiers revealed more distinct differences to the ensembles (see Fig. 3 and Table 1 and 2).

**Figure 3.**
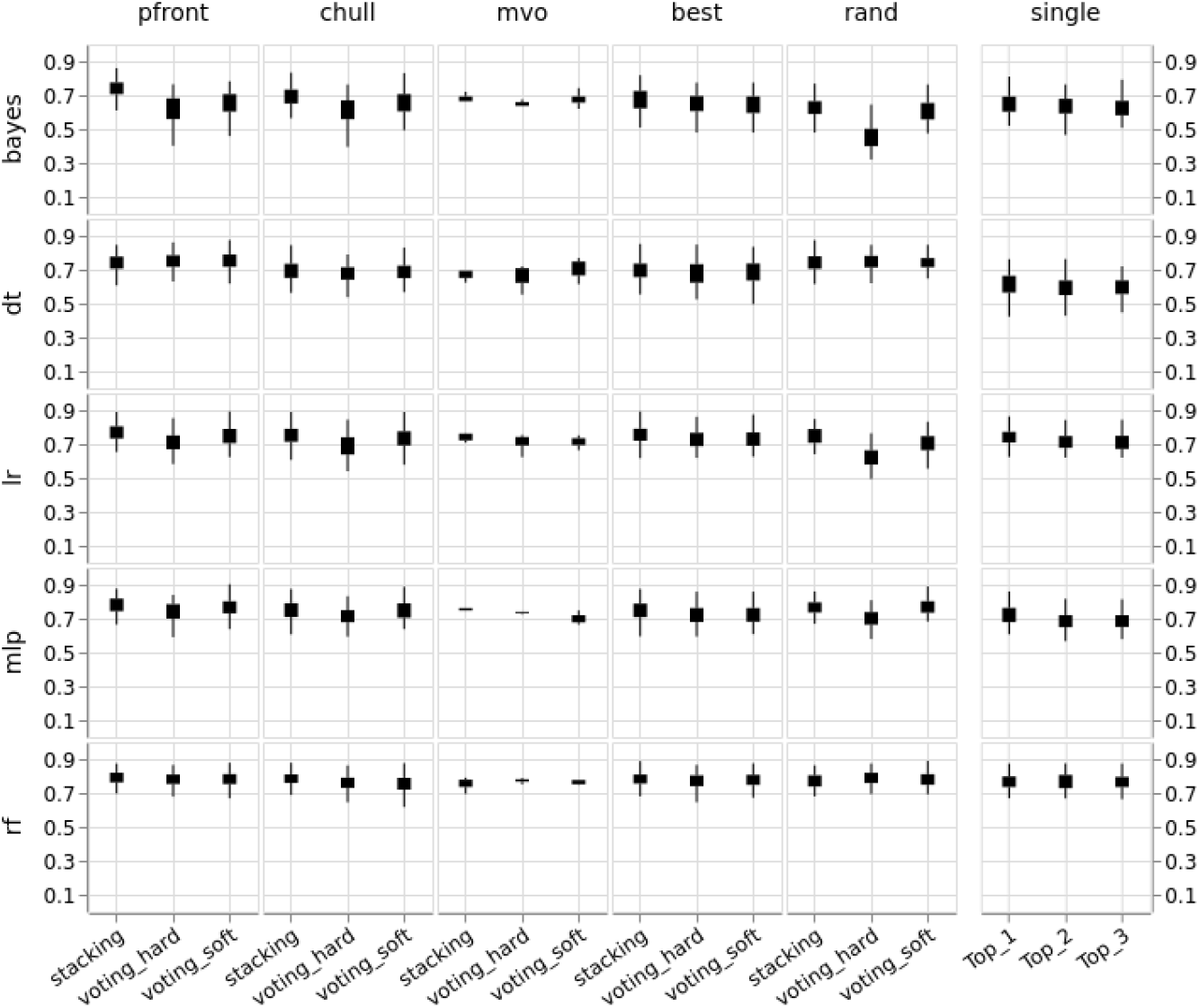
The box plot shows the Matthews correlation coefficients (MCC) distribution (y-axis) using a 100-fold Monte Carlo cross-validation. The rows depict the base with various encodings, and the columns depict the pruning method. The respective ensemble types can be found in the x-axis. The MVO pruning (mvo) has only been conducted on the first five folds (see Discussion). The MCC of the individual classifiers (single) has been collected for the best (Top 1), second (Top 2), and third (Top 3) model per fold, respectively. The plot shows the example of the avp amppred dataset.

### Data visualization

We leveraged two standard visualization techniques, which we adapted and extended for our particular application. First, we enhanced the kappa-error diagram [25] for the presentation of multiple folds, i.e., 100 in the current study, by aggregating the cross-validation results into a two-dimensional histogram (see Fig. 4). The color code allows the viewer to spot the peak at one glance. Hence, the tendency of ensembles to use a specific base classifier. Moreover, considering the distribution of the variables, one can make conclusions about the robustness.

**Figure 4.**
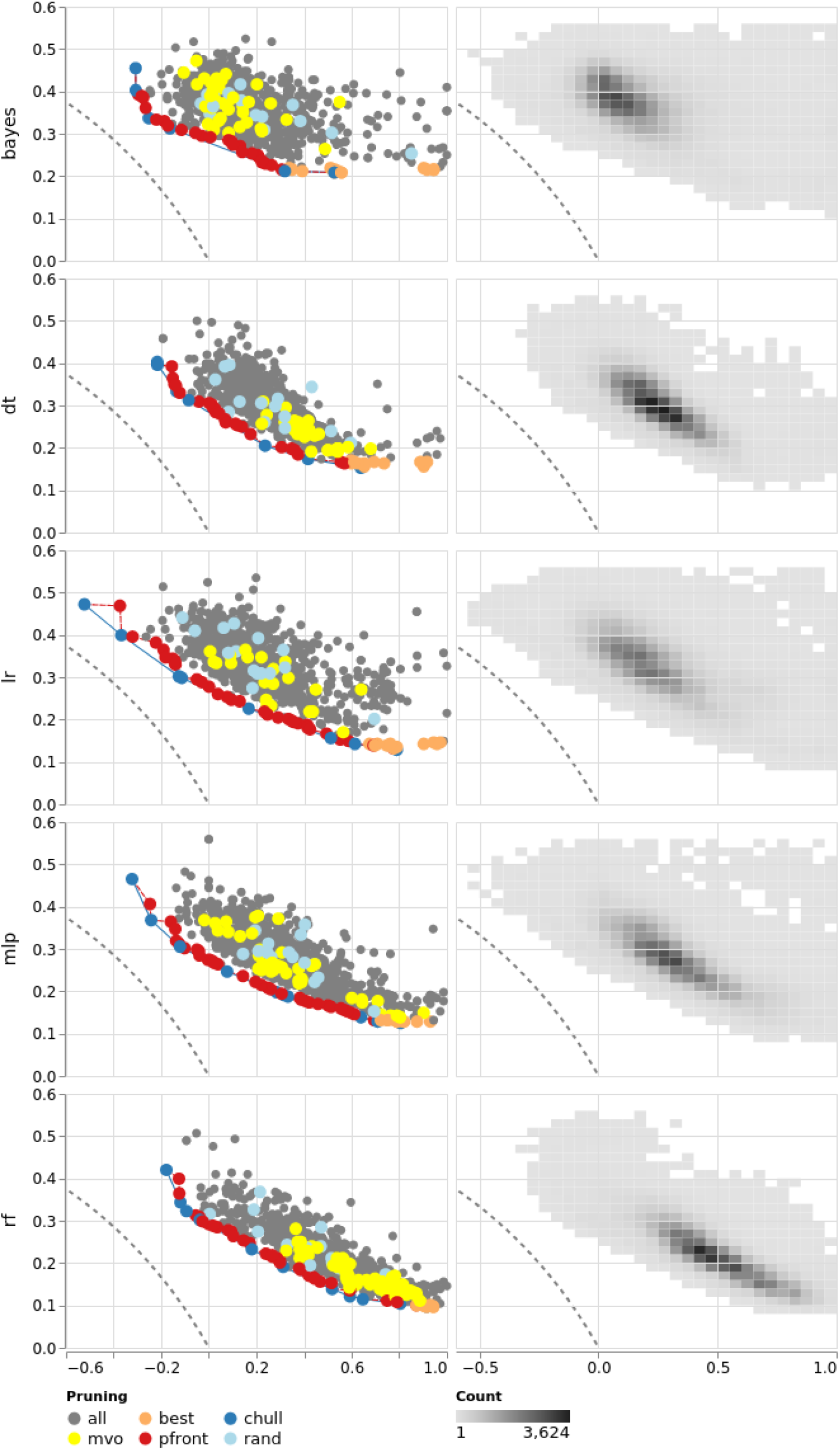
The kappa-error diagram depicts pairs of base classifiers (dots) using the kappa diversity on the x-axis and the average error on the y-axis (left column). Each classifier pair denotes two particular encodings. Rows depict the base classifiers. The dark blue line indicates the convex hull, and the red line the Pareto frontier. Pruning methods are represented as colors. The left column shows the first fold. The right column groups the results of all folds in a 2D histogram. The darker the color, the more classifier pairs are binned in one group. The dashed line in all panels depicts the theoretical boundary (refer to the method section for more details). The plot shows the example of the avp amppred dataset.

Second, we extended the critical difference (CD) chart [35] with a categorical heatmap displaying the actual performance. The extension enables viewers to compare classifiers and review the individual encoding performance, i.e., Matthews correlation coefficient in the present case, at one glance. In addition, the thickness of the vertical and horizontal rules is directly related to the critical difference, i.e., the thicker the rule, the closer the classifiers to the critical difference. Thus, the rule thickness provides an additional visual channel to access the CD.

## Discussion

We developed a workflow for unsupervised encoding selection and performance assessment of multiple ensembles and base classifiers. Thus, we implemented and compared several algorithms to facilitate ensemble pruning, including convex hull, Pareto frontier pruning, and multi-verse optimization (MVO). Our results demonstrate that the crucial factors are the base classifiers and the individual encodings. The ensemble technique was not relevant, i.e., we could not observe performance variations using one of hard or soft voting or stacking. In general, applying the Random Forest (RF) or Multi-Layer Perceptron (MLP) as a base classifier yielded good performance across all datasets. The Pareto frontier pruning selected suitable encodings throughout the experiments. In addition, we observed similar performance as reported in the original studies (see Tables 1 and 5).

**Table 5.**
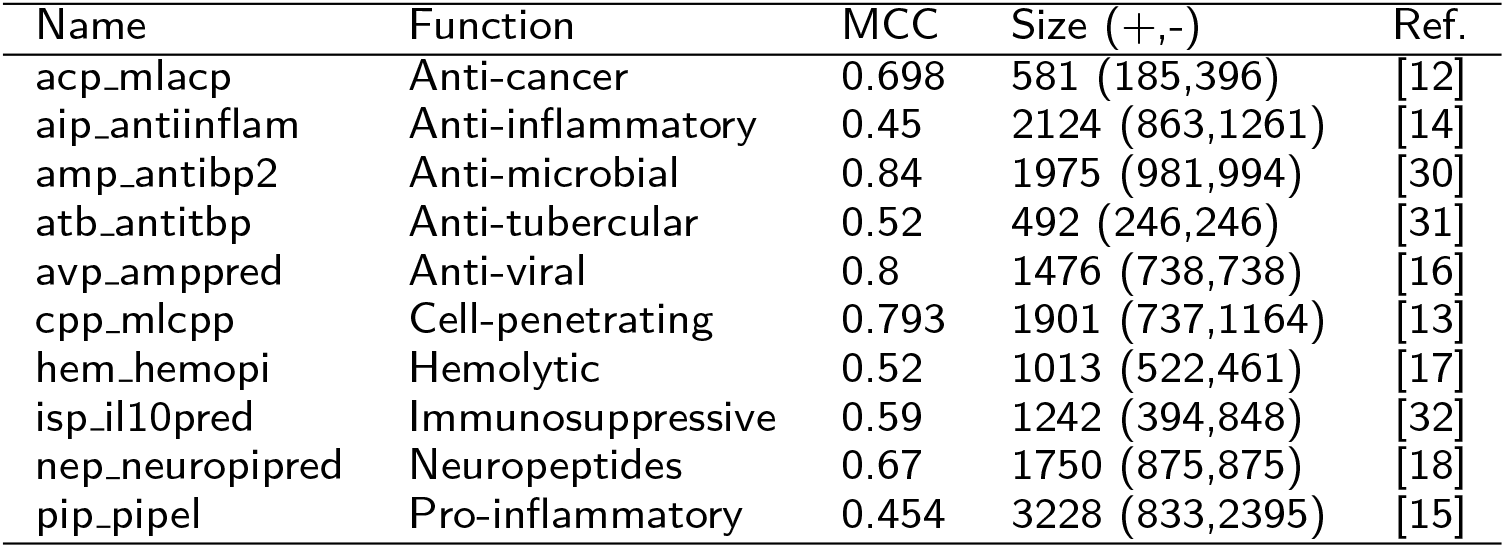
Employed datasets in this study. The function refers to the positive class, i.e., sequences of class + possess the respective function. The stated MCC refers to the performance reported in the original study. See the references or [20] for more details.

However, since we used one encoding per base classifier, we restricted the employed ensemble methods, i.e., majority voting, averaging, and stacking, which do not modify the base classifiers. These ensemble types are in contrast to others, e.g., boosting, where weights are adapted for misclassified training instances in base classifiers [36]. More research is necessary to investigate how performance and more sophisticated ensemble methods are associated. The employed ensemble types are also the reason for the kappa-error point cloud shape solely depending on the base classifiers. Consequently, computing the kappa-error diagram for all ensemble methods was not necessary. Our encoding/classifier approach is also contrary to other studies, e.g., [12], [14], or [16], which concatenated several encoded datasets to one final dataset (hybrid model) and applied feature selection before training. In the present study, we solely scaled the datasets to standardize the feature range, and used the encoded datasets largely unprocessed, potentially affecting the final performance. Furthermore, we neglected hyperparameter optimization of the base classifiers and focused on encoding, pruning and model selection. In this respect, more research is necessary to investigate the impact of hyperparameter optimization of the base models on the ensemble performance. However, since our tool determines important components of effective ensembles, researchers can subsequently improve the hyperparamters, if necessary.

In general, utilizing the Pareto frontier pruning generates good ensembles; however, requiring the calculation of the Cartesian product of all base classifiers; thus, encodings. Although only the (lower) triangular matrix is necessary, the computation is still CPU-intensive.

Another point of concern refers to the selection of exactly 15 classifier pairs per fold for the random and best encoding ensembles. In contrast, the size of classifier the Pareto frontier and convex hull pruning ensembles depend on the number of classifier pairs on the Pareto front and convex hull, respectively. Thus, a fair comparison is difficult. However, we repeated the experiments multiple times using a 100-fold MCCV, which rules out any selection bias. Considering the performance between individual models and ensembles, the contribution of the diversity is only slight, however, more research is required in this direction.

The results of the MVO also acknowledge the impact of diversity. One can observe that the MVO generates inferior ensembles (see Fig. 3). However, due to the limited number of generations, the MVO could be stuck in local optimum (see Fig. 1). More research is also required to examine specific advantages and disadvantages of the MVO for encoding selection [37].

In the kappa-error chart, which depicts preferable classifier pairs towards the lower-left corner, one can readily recognize the inferiority of the MVO (see Fig. 4). The classifier pairs are distributed across the kappa-error area, i.e., the MVO screens the entire solution space and adds weak classifiers to the final ensemble. Nevertheless, since we limited the maximum number of generations to 15, we cannot rule out that more generations would yield better results. Moreover, due to high resource consumption, we limited the MVO to 5 folds, which hampers the comparison.

Moreover, the Random Forest (RF) deployment as a single classifier reveals good performance, which is not surprising since it is already an ensemble algorithm. In this respect, the other base classifiers are less accurate (see Fig. 3). However, it could be demonstrated that RFs as base classifiers, i.e., using different encoded datasets per model, slightly improves the performance. This further highlights the importance of different encodings, hence the projection of different biological aspects, for the classification process.

The implemented methods demonstrate usability on a broad range of datasets from various biomedical domains. We incorporated the MVO owing to its good performance on several benchmark datasets [38]. The comprehensive Monte Carlo cross-validation copes with the variance, ultimately increasing the robustness of the results. In addition, the Pareto frontier and convex hull pruning consider simultaneously the performance and the diversity of encodings and base classifiers; hence, compensating their strength and weaknesses and revealing their potential for ensembles [39]. Our proposed extension to the critical difference chart allows the viewer at one glance to grasp significant, i.e., critical, performance differences of encodings, models, and pruning methods jointly with the actual performance.

## Conclusions

In summary, we employed two overproduce-and-select methods, namely Pareto frontier and convex hull pruning, as well as the multi-verse optimizer for exhaustively searching the encoding/base classifier space. We employed Logistic Regression, Decision Trees, Näive Bayes, Multi-Layer Perceptron, and Random Forest as base models and majority vote, averaging, and stacked generalization for the fusion. The experiments and visualizations enable the comparison of the respective components; however, further research is necessary to examine other ensemble classifiers, e.g., boosting. We considered ten datasets as case studies.

All in all, we propose an extensible workflow for automated encoding selection through diverse ensemble pruning methods. Depending on the use-case, i.e., the initial encoding space, our workflow can be used stand-alone, or as an extension for our recent work [20], ultimately easing the access for non-technical users. In case of the avp amppred dataset, we observed good performance using the Pareto frontier pruning. Our workflow determines the best encodings, base models, as well as pruning, and fusion method for the classification task at hand. This allows researchers to built upon an effective model and focus on hyperparameter optimization afterward.

## Methods

We developed a workflow using Snakemake v6.5.1 [40], Python v3.9.1, and R v4.1.0. For the machine learning algorithms, we employed scikit-learn v0.24.2 [41]. We will use the following definitions throughout the manuscript: the original unprocessed dataset is denoted as the dataset. One dataset can be encoded in manifold ways, which we refer to as encoded datasets. Finallly, encodings specify particular encoding algorithms.

The peptide datasets are taken from the PEPTIDE REACToR [20]. We also utilized this tool to reduce the initial set of encoded datasets. Ultimately, the goal of our workflow is the selection of the best encodings from the reduced set of encoded datasets. Note that best encodings refers to datasets, encoded with specific, i.e., effective, encoding algorithms. Afterward, the encoded datasets are used to train the base classifiers.

There are two approaches to harness multiple encodings in a single model, namely the fusion and the hybrid model [21]. Fusion models train one encoding per base classifier and fuse the output for the final prediction. Contrary, hybrid models use the concatenated features of multiple encodings for single model training. The concatenation approach is particularly problematic for entropy-based models such as DT or RF due to the bias in variable selection. Thus, in the present study, we implemented the fusion design, i.e., each ensemble consists of an arbitrary amount of base classifiers using one particular encoding, respectively. Finally, the employed datasets from a wide range of biomedical domains ensure broad applicability and the robustness of our results.

The workflow conducts the following steps. First, train, validation, and test indices are calculated to ensure equal samples for the cross-validation. More precisely, we computed the indices before the experiments such that all base and ensemble models use the same train, validation, and test data in each fold. Second, we standardized the encoded datasets using a min-max normalization between 0 and 1. Afterward, we trained and assessed models for all encoded datasets and ensemble types using a 100-fold Monte Carlo cross-validation. Besides the pruning algorithms, we selected the best and random encodings and compare the results to individual models. Finally, we statistically assessed and visualized the results (see Fig. 5). Significant steps are described in more detail below.

**Figure 5.**
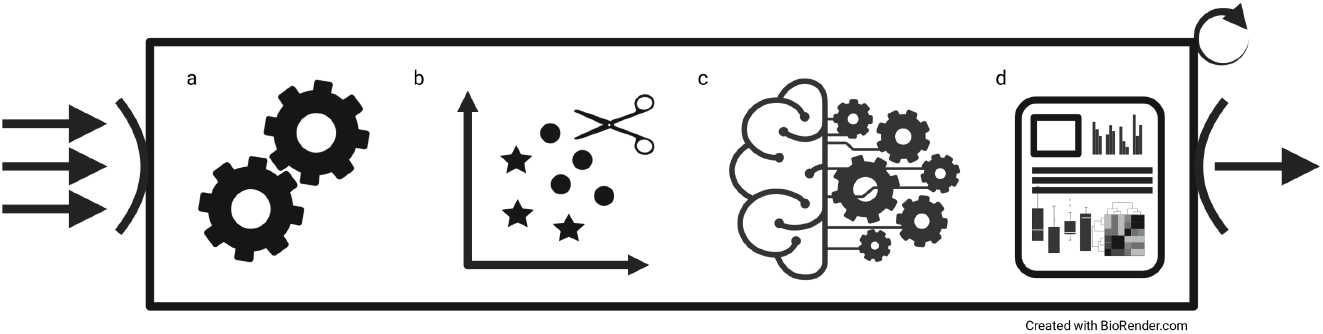
Overview of the workflow. (**a**) For each fold of the MCCV, the preprocessing is conducted, i.e., the indices of the train/test splits are determined, and the data is scaled. (**b**) The pruning methods, e.g., Pareto frontier and MVO, select the current fold’s encodings and the number of base classifiers. (**c**) Different ensembles with various base classifiers are trained and validated on the test data. (**d**) The results are collected, statistically validated, and illustrated. The workflow accepts an arbitrary number of datasets as input (arrows). For each dataset (bold arrows), the steps a to d are executed successively. Refer to the method section for more details.

Note that we used Matthews correlation coefficient (MCC) throughout the study to handle the imbalance in the datasets [42]:

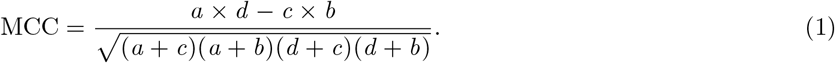

*a* is the number of true positives, *d* is the number of true negatives, *b* is the number of false negatives, and *c* is the number of false positives.

### Datasets

For a comprehensive analysis on peptide encodings, Spänig *et al*. (2021) gathered and encoded a variety of datasets from multiple biomedical domains [20]. Details about the dataset preprocessing and encoding algorithms can be found in this publication [20]. We selected datasets with low to medium classification performance from this collection, i.e., a reported MCC of 0.63 *±* 0.15 on the independent test set; additionally, covering diverse biomedical applications. Moreover, we excluded datasets for which accurate models have been published to investigate the potential effects of different classifiers and ensembles. We limited our study to ten datasets to cope with the computational complexity. The dataset size ranges from 492 to 3,228 sequences with an average of 1, 580.8 *±* 812.1 sequences. The datasets comprise 15,782 sequences with a mean length of 21.17 *±* 13.23 amino acids. 6,404 sequences belong to the positive and 9,378 to the negative class. The average sequence length is 22.47 *±* 15.88 and 20.29 *±* 10.97, respectively. Duplicated sequences have been removed. Refer to Table 5 for more details.

### Monte Carlo cross-validation

We applied a 100-fold Monte Carlo cross-validation (MCCV) [43]. The MCCV improves the generalization and diminishes the variance of the results, i.e., results are more robust, hence comparable. In addition, we ensured that the *n*-th fold is identical across all experiments leading to improved comparability across all base classifiers and ensembles. Each fold is composed of one split using 80 % of the data for training, 10 % for validation, and the remaining 10 % for testing (see Fig. 6). In contrast to *k*-fold cross-validation, MCCV follows a sampling with replacement strategy, i.e., splits can contain identical samples multiple times. However, duplicate samples do not occur in the train, validation, and the test split [43].

**Figure 6.**
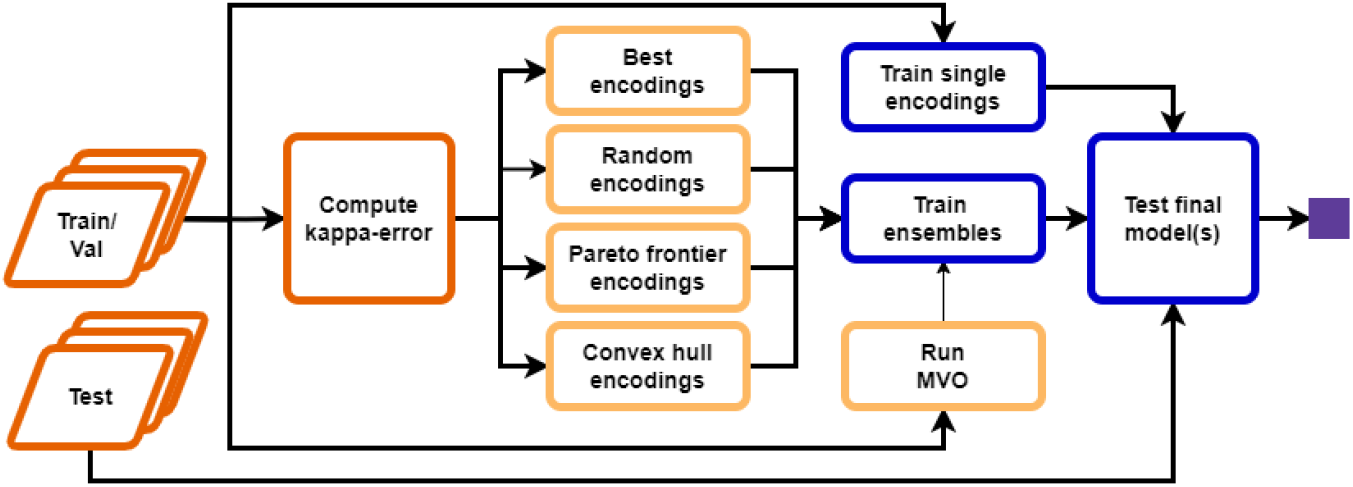
Overview of the cross validation. (**Orange**) In the preprocessing the train/test splits are created, and the kappa-error diagram is calculated. (**Yellow**) The pruning methods, including MVO, select the best encodings and base classifiers based on the validation data. (**Blue**) All models are trained with the entire training data and tested on the unseen test set. (**Purple**) The results are collected, statistically validated, and illustrated.

### Base classifiers

We used the following base classifiers for our experiments: Näive Bayes, Logistic Regression, Decision Tree, Multi-Layer Perceptron, and Random Forest. Each classifier will be briefly described hereinafter. We used the implementations provided by the scikit-learn library [41]. We utilized the default hyperparameters.

### Näive Bayes

The Näive Bayes (NB) classifier (naively) assumes conditional independence of the feature vectors and applies the Bayes theorem for the prediction [25]. Model training is enabled via a probability density function (PDF) and the prior probability of a given class. For simplicity, we assume a Gaussian distribution of the features. Hence, we applied the Gaussian NB using

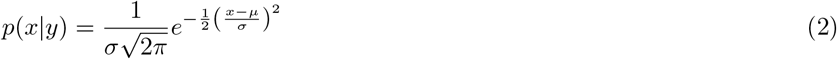

as the PDF, whereby *σ* denotes the standard deviation and *µ* the mean of features *x* given a class *y* [44].

### Logistic Regression

The binary Logistic Regression (LR) is another probability-based classifier, i.e., it derives the probability of a class *y* given a feature vector *x* [45]. The LR predicts probabilities between 0 and 1 using the logistic function denoted as

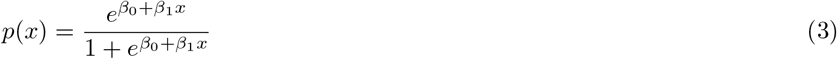

and the maximum likelihood function to estimate the coefficients *β*, i.e., to train the model [45].

### Decision Tree

The Decision Tree (DT) classifier, precisely the CART (Classification And Regression Trees) implementation, is a tree-based model, i.e., a tree structure is generated during training [46]. Each node is based on the most discriminating feature [25]. New splits are created based on the Gini impurity *i* of the remaining data:

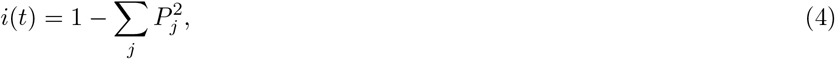

where *j ∈ {*0, 1*}* for binary classification and *P* is the probability of class *j* at a node *t* [25]. If a split is pure enough, a leaf node is added. Otherwise, intermediate nodes are created [25]. For prediction, the tree is traced until a leaf node, which states the final class.

### Multi-Layer Perceptron

The Multi-Layer Perceptron (MLP) follows the functioning of neural networks in the human brain [25]. In particular, the MLP is an artificial neural network consisting of multiple layers of neurons, which are connected with a certain intensity, i.e., are weighted. Based on the input and the activation function of the neurons, the MLP propagate the input data forward and finally assigns the class in the last layer. To train the MLP, the data is backward propagated and the weights are updated accordingly to minimize the training error [25].

### Random Forest

The Random Forest (RF) classifier is an ensemble learning technique, which trains multiple DTs on random samples, i.e., bagging, of the input data [47]. For the final classification, the majority vote of the trees is used [47]. Note that we use the RF as a base learner, which allows comparing the performance with DTs and the actual ensembles techniques in general (see below).

### Classifier ensembles

To combine the output of the base classifiers introduced above, we employed the following ensemble methods: majority vote (hard voting), averaging (soft voting), and stacked generalization (stacking). In the present study, each base classifier is trained on one encoded dataset, meaning if for one dataset *n* encodings are selected, the size of one ensemble is *n*. We adapted the implementations of the scikit-learn library [41], such that not only one dataset but several encoded datasets can be used for training. For instance, if one passes *n* encoded datasets, the ensemble consists of *n* base classifiers trained on one particular encoded dataset, respectively.

#### Majority voting

The majority voting ensemble (hard voting) combines the output by ultimately assigning the class, which has been predicted by the majority of the single base classifiers. We employed the customized version of scikit-learn’s VotingClassifier class with hard voting enabled.

#### Averaging

The averaging method (soft voting) computes the means of the predicted class probabilities per base classifier. The maximum value determines the final class. We used the adjusted VotingClassifier with voting set to soft.

#### Stacked generalization

The stacking approach utilizes the output of the base classifiers to train a meta-model, i.e., the predicted class probabilities of the base classifiers are used as features [48]. We adapted the StackingClassifier from the scikit-learn package and employed Logistic Regression as the meta-model.

### Ensemble pruning

Selecting the correct number of base classifiers in an ensemble is challenging. Thus, Kuncheva (2014) suggests several approaches to determine the ensemble size [25]. For instance, sequential forward selection, adding one classifier successively, in case the additional model improves the ensemble performance [25]. However, in the present case, we are dealing with potentially hundreds of encoded datasets, for which this particular technique is not practical. To this end, we used two selection methods, namely convex hull and Pareto frontier pruning, which handle the complexity better [25]. Moreover, we implemented the multi-verse optimization algorithm as an additional encoding selection technique [49]. Finally, we employed best and random encodings selection as a reference. The pruning methods are described more detailed in the following.

### Kappa-error diagram

The kappa-error diagram, introduced by Margineantu and Dietterich (1997), is the basis for the convex hull and Pareto frontier pruning [50]. The graph represents pairs of classifiers by their average error and diversity, as shown in Fig. 4. The diversity measures the agreement of classifier outputs, i.e., the better the agreement of the classifier predictions, the less the diversity [25]. Specifically, the kappa diversity is denoted as

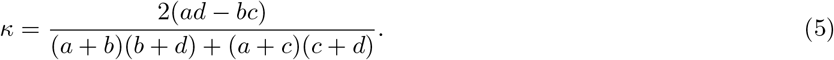

The *κ* statistic ranges from *−*1 to 1, whereby *κ* = 1 denotes perfect agreement, *κ* = 0 random, and *κ <* 0 worse than random consensus [50]. The error is calculated using

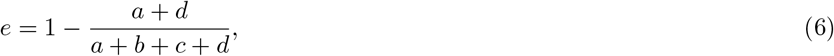

with the subtrahend being the accuracy. However, Kuncheva (2013) pointed out that diversity concerning the average error can not be arbitrarily low [39]. In fact, desirable classifier pairs approximate the lower-left corner (see Fig. 4), i.e., approximating a theoretical boundary, which is defined in Eq. 7 [39].

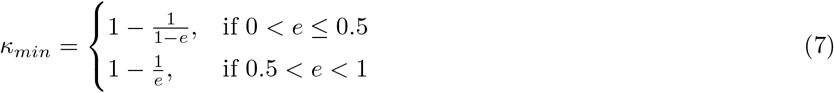

Note that the classifier pairs are composed using the lower triangular matrix of the Cartesian product. Afterward, the pruning methods select a subset of pairs, also likely include duplicated base classifiers. Thus, all pruning methods ensure that the final ensemble only uses unique classifiers. Hence, base classifiers are trained on individual encoded datasets.

### Convex hull

The kappa-error diagram depicts a set of points, i.e., pairs of base classifiers, in a two-dimensional space. The kappa diversity is the first, and the pairwise average error is the second dimension. We employed the Quickhull algorithm to calculate the convex hull [51]. Hence, the smallest convex set that contains the classifier pairs [51]. Thus, no further classifier pairs exist beyond the convex hull. We utilized the implementation of the Quickhull algorithm provided by the SciPy package in the ConvexHull module [52].

Since we are only interested in the partial convex hull, that is, pairs approaching the theoretical boundary defined in Eq. 7 and depicted in Fig. 4, we adapted the *pareto n* algorithm from Kuncheva (2014), which returns only classifier pairs fulfilling the criteria [25].

### Pareto frontier

The Pareto optimality describes the compromise of multiple properties towards optimizing a single objective [53]. For instance, a pair of classifiers is Pareto optimal if improving the diversity is impossible without simultaneously impairing the average pairwise error. Analog to the partial convex hull introduced earlier, Pareto optimal classifier pairs approach the theoretical boundary as stated in Eq. 7, ultimately defining the Pareto frontier. Again, we used the *pareto n* algorithm adapted from Kuncheva (2014) to obtain all classifier pairs determining the Pareto frontier (see Fig. 4).

### Multi-verse optimization

The multi-verse optimization (MVO) algorithm is inspired by the alternative cosmological model stating that several big bangs created multiple, parallel existing universes, which are connected by black and white holes and wormholes [38]. In terms of an optimization algorithm, black and white holes are used to explore the search space and wormholes to refine solutions [38]. Moreover, the inflation rate, i.e., the fitness, of universes is used for the emergence of new holes; thus, to cope with local minima [49]. We selected the MVO owing to its superior performance compared to other state-af-the-art optimization algorithms, for instance, Particle Swarm Optimization [49]. An outstanding property of the MVO study is the efficiency of searching and optimization [49]. This property is beneficial for the selection process, since the parameterized encoding groups form a large search space [20]. For more details, refer to Mirjalili *et al*. (2016) and Al-Madi *et al*. (2019) [38, 49].

We implemented the binary MVO following [49] using Python. Each solution candidate is represented as a binary vector, where each position denotes the path to an encoded dataset, that is, the *i*-th bit set means that the *i*-th encoding is included in the final ensemble (see Fig. 4). We examined different generations, i.e., 100, 80, 50, 25, and 15. However, we observed that performance depends mainly on the initialization and count of the universes. Specifically, the performance gain from the 15th generation is minor but requires much time. Thus, we set the optimization to a maximum of 15 generations with 32 universes each. Due to its resource intensity, we executed the MVO only for the first five folds (see section Monte Carlo crossvalidation).

### Best encodings

A further pruning method uses only the best classifier pairs. In particular, based on the kappa-error diagram, the algorithm selects 15 classifier pairs with the lowest pairwise average error (see Fig. 4).

### Random encodings

The last pruning method selects 15 random classifier pairs from the kappa-error diagram. Note that the selection is only performed one time. That is, the pairs are the same across all folds.

### Statistics

We examined the areas covered by the respective base classifiers (see Fig. 4). To this end, we calculated the area for each fold. The area is described by multiple variables, i.e., the kappa diversity and the average pairwise error. Thus, we applied the multivariate analysis of variance (MANOVA) to verify if the areas differ significantly. If this is the case, we subsequently employed an analysis of variance (ANOVA) to investigate the effect of the diversity and the average error separated. For post-hoc assessment, Tukey’s HSD has been applied. We used the tests provided by the R standard library. *α* was set to 0.05, i.e., p values *≤* 0.05 are considered as significant.

In addition, we employed the Friedman test with the Iman and Davenport correction for the statistical comparison of multiple single and ensemble classifiers [54]. In the case at least one model is significantly different, we used the Nemenyi test for post-hoc analysis [54]. Refer also to Spänig *et al*. (2021) for more details [20]. The tests were provided by the scmamp R package v0.2.55 [35].

Finally, we examined if the best ensemble has a significant improvement over the best single classifier using Student’s *t* -test for repeated measures, i.e., paired samples. Again, *α* was defined as 0.05.

### Data visualization

All plots are realized using Altair v4.1.0 [55] and described in more detail hereinafter.

#### Kappa-error diagram

The kappa-error diagram, suggested by Margineantu and Dietterich (1997) [50], shows the result of a single split in the left column and a two-dimensional histogram aggregating all folds in the right column (see Fig. 4). The rows show the base classifiers. Note that the kappa-error shape depends only on the base classifiers (see Discussion). The left column also visualizes the partial convex hull (black line) and the Pareto frontier (red line). The colors refer to the pruning method. Each dot is a classifier pair trained on two encoded datasets. Note that we display only 1000 dots per panel. Moreover, we set the bin size to 40 for the binned heatmap with darker colors depicting more values.

#### XCD chart

The extended critical difference (XCD) chart (Fig. 2) is based on the critical difference chart introduced by Calvo and Santafé (2016) [35]. Classifier groups not surpassing the critical difference (CD) are connected with black lines. The line thickness depicts the actual CD, meaning groups associated with thicker lines are closer to CD. The XCD charts present two classifier groups. The x-axis includes pruning types, and the y-axis the actual ensembles and the corresponding base classifier. The main area contains a categorical heatmap showing Matthews correlation coefficient (MCC) in 0.05 steps. The darker, the higher the MCC. The MCC is the median MCC of the respective group combination and corresponds to the median from Fig. 3. Note that for the computation of the CD, we concatenated the MCCs of all cross-validation runs, e.g., 12 * 100 MCCs for pfront, and 6 * 100 MCCs for bayes voting soft.

## Supporting information

Supplement 1 (interactive plots)

Supplement 2 (dataset results)

## Availability of data and materials

The source code can be found at https://github.com/spaenigs/ensemble-performance. All datasets are available at https://github.com/spaenigs/peptidereactor.

## Ethics approval and consent to participate

Not applicable.

## Consent for publication

Not applicable.

## Authors’ contributions

SS and DH developed the concept. SS designed and performed the experiments and analyzed the data. SS and DH interpreted the results. AM implemented the MVO algorithm. SS wrote the manuscript. DH supervised the study and revised the manuscript. All authors read and approved the final manuscript.

